# Region-specific magnetic fields structure sea turtle populations

**DOI:** 10.1101/630152

**Authors:** Sahmorie J.K. Cameron, Miguel Baltazar-Soares, Christophe Eizaguirre

**Affiliations:** Queen Mary University of London, School of Biological and Chemical Sciences, Mile End Road, E14NS, London, United Kingdom

## Abstract

Philopatry and long distance migrations are common in the animal kingdom, of which sea turtles are flagship examples. Recent studies have suggested sea turtles use the Earth’s magnetic field to navigate across ocean basins to return to their natal area, yet the mechanisms underlying this process remain unknown. If true though, the genetic structure at nesting sites should positively correlate with differences in location-specific magnetic vectors within nesting regions. Here, we confirm this working hypothesis but only in certain regions of the world and for all sea turtle species nesting in those regions. Reversely, where no correlations were found between genetic differentiation and geomagnetic vectors, this was the case for all nesting sea turtle species. Our approach hence reveals parallel but not universal use of geomagnetic cues in sea turtles. We describe magneto-sensing regions as characterized by sharp clines of total and vertical field intensity vectors offering the navigation cues that increase philopatric accuracy and promote genetic structuring among sea turtle populations.

## Introduction

Philopatry is a well-documented trait of many species, including insects, birds, mammals and reptiles (Greenwood 1980; FitzSimmons *et al*. 1997). With this behavior, after long-distance migration, individuals return to their natal area upon sexual maturity to mate, sometimes decades after leaving at an early life stages. While philopatry can be a learned trait from older individuals (e.g. sheep, moose (Jesmer *et al*. 2018), whales (Baker *et al*. 2013)), in others, it is an evolved behaviour associated with long-distance navigation and imprinting, such as butterflies, salmon, and sea turtles (Brower 1995; Dittman & Quinn 1996; Lohmann *et al*. 1999). Sea turtles provide perhaps the best illustration of philopatry in the marine environment since they return to their natal area after ~25-30 years and thousands of kms of juvenile migration (Wyneken *et al*. 2013). Philopatry can be as accurate as 50 km (Hatase *et al*. 2002; Bowen & Karl 2007) and shapes population structure (Stiebens *et al*. 2013b). Yet, its underlying mechanisms remain partly speculative.

Several studies have demonstrated that the Earth’s magnetic field provides sufficient cues to guide animal migration (Wiltschko & Wiltschko 2005; Lohmann *et al*. 2008b). The Earth magnetism is driven by the motion of the molten iron in Earth’s core and is defined by seven vectors (Campbell 2003). The shifting of liquid iron also means the magnetic field is in a constant state of flux over various time-scales. While it is acknowledged that imprinting occurs during the early stages of development or birth (Lohmann *et al*. 2008b), the underlying mechanisms of geomagnetic navigation remain unknown, particularly in sea turtles. Nevertheless, if, globally, geomagnetic imprinting determines philopatric behaviour (Brothers & Lohmann 2015), it should leave a signature of selection on the population genetic structure of sea turtles. This is because, within a region, if a location’s (nesting site or natal area) geomagnetic profile differs extensively from neighbouring locations, the navigation accuracy of sea turtles should be high and genetic structure should increase, i.e. gene flow among populations should decrease.

By examining the possible determinants of genetic structure across the nesting aggregations (regions) of sea turtles, and across species with overlapping nesting distributions, we should be able to disentangle species-specific features of magneto-sensing from region-specific geomagnetic characteristics. This approach compensates for the lack of known underlying molecular mechanisms and the associated cline analysis of allele frequency change (Günther & Coop 2013). Positive correlations between geomagnetic field vectors and population genetic structure would confirm that sea turtles use vectors of the earth magnetic field for their philopatric migration. Finding no such pattern would challenge the imprinting hypothesis of early life stages as a potentially universal mechanism of navigation.

Here, we combine a thorough literature search with sampling and mitochondrial control region sequencing. We were able to obtain population structure data from five of the seven sea turtle species from 144 locations (nesting sites and natal areas) in nine regions globally, for ~ 17,470 sequences (Fig. 1). We obtained data from nesting regions where several species co-exist allowing us to disentangle species from region-specific effects and explore whether all species are equally responsive to geomagnetism. Specifically, we tested the correlation between genetic structure, estimated from Wright fixation indices, and geomagnetic vectors at natal areas. F_ST_ fixation indices from mitochondrial genes are widely used as markers of philopatric behaviours because of their maternal mode of inheritance (Bowen & Karl 2007).

**Fig. 1:**
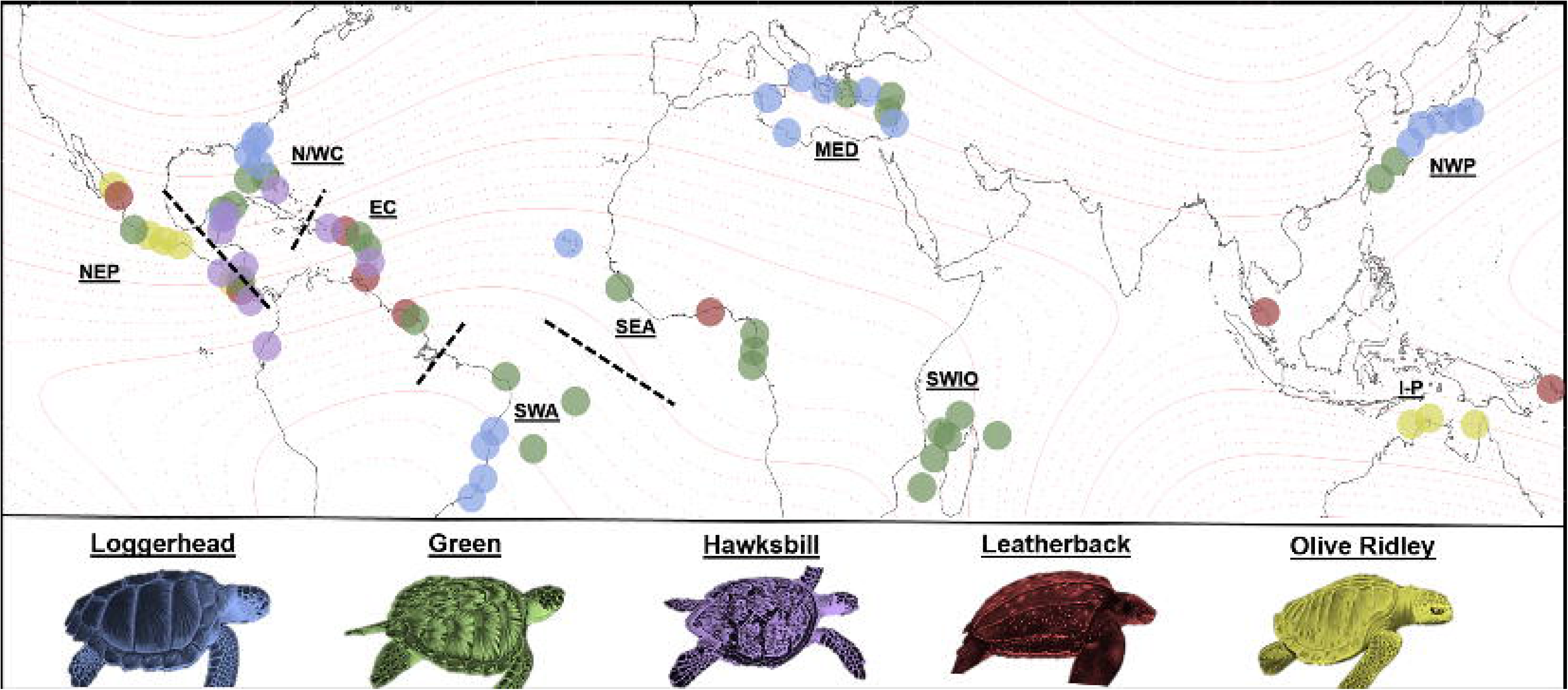
World Magnetic Model (Epoch 2015.0) of total field intensity. Turtle species, locations split by nesting regions and field intensity isolines (□=1000nT) are shown. Abbreviated nesting regions: EC = Eastern Caribbean, I-P = Indo-Pacific, MED = Mediterranean, NEP = Northeast Pacific, N/WC = Northern/Western Caribbean, NWP = Northwest Pacific, SEA = Southeast Atlantic, SWA = Southwest Atlantic, SWIO = Southwest Indian Ocean.

## Methods

### Literature search and data collection

We took advantage of the well-characterized population genetic structure of sea turtle across the globe. Data were collected from published literature following a two-step process. Firstly, we used SWOT (The State of the World’s Sea Turtles), IUCN (International Union for Conservation of Nature) and the grey literature to identify all the world nesting aggregations (Electronic supplementary material, Table S1). Secondly, we screened the primary and secondary literature for all studies with indices of population structure expressed as a pairwise Wright’s fixation index (F_ST_, γ_ST_ or φ_ST_). To find this literature, we searched for “Sea turtle AND” either “Phylogeography”, “Genetic structure”, “Population structure” or “Mitochondrial DNA” together with the names of the nesting aggregations. We also performed individual searches for each of the seven sea turtle species substituting “Sea turtle” for: “Loggerhead”, “Green turtle”, “Hawksbill”, “Leatherback”, “Flatback”, “Kemp’s Ridley” and “Olive Ridley”.

As our goal was to specifically compare genetic and geomagnetic information within sea turtle rookeries, obtaining exact nesting beach locations to derive accurate magnetic field information was important. Therefore, if the literature contained only one nesting site from a specific nesting region, this data point was removed (5 nesting sites removed). For literature that contained both nesting site and feeding ground genetic information, feeding ground data points were ignored. Data points in which genetic structure information was pooled from multiple geographic locations into a single general natal area were removed (18 nesting sites removed). When the exact nesting site sampled for a location could not be identified, data points were also removed (3 nesting sites removed). This resulted in the use of 144 locations (Electronic supplementary material, Table S2).

Nesting sites and areas among sea turtle species were grouped into what we define as “nesting regions” based on geographic location. We did not use regional management units (RMUs) due to the lack of consistency of RMUs among species, which would prevent direct comparison. It resulted into a dataset composed of 144 locations, from 9 different nesting regions worldwide for 5 of the 7 sea turtle species. Noteworthy, as gene flow across regions does not occur, data points between nesting regions did not enter the analyses to avoid biasing for high values of genetic structure indices.

### Population structure in the largest North East Atlantic loggerhead turtle rookery

To complement the existing information, we also added data from the Cape Verde Archipelago, the 3^rd^ largest aggregation of loggerhead turtles in the world. Field surveys took place in the Cape Verde archipelago in 2011, 2012 and 2013 during the nesting season from June to October. Nine different islands were sampled: Boavista, Fogo, Maio, Sal, Santa Luzia, Santiago, Santo Antão, São Nicolau and São Vicente. A 3mm piece of non-keratinized tissue was removed from the right front flipper using a single-use disposable scalpel and samples were stored in ethanol (Stiebens *et al*. 2013a; Stiebens *et al*. 2013b). Turtles were tagged with metal tags and/or pit tags directly after egg deposition to track nesting behaviour and to avoid multiple sampling. In the laboratory, each sample was washed in distilled water for 20 seconds and cut into smaller pieces. DNA from tissues was extracted using the DNeasy^®^ 96 Blood & Tissue Kit (QIAGEN, Hilden, Germany). Elution was conducted in twice 75 μl of AE Buffer. All other steps followed the manufacturer’s protocol.

The control region of the mitochondrial DNA was amplified using the Primers LCM15382 (5’-GCTTAACCCTAAAGCATTGG-3‘) and H950 (5’-GTCTCGGATTTAGGGGTTTG-3‘)(Abreu-Grobois *et al*. 2006) following (Stiebens *et al*. 2013a). PCR products were cleaned with ExoSAP-IT^®^ following the manufacturer’s protocol. Cycle sequencing reactions were performed with Big Dye^®^ Terminator v3.1 Cycle Sequencing Kit (Applied Biosystems, Darmstadt, Germany). Sequences were obtained from the forward direction. In cases where a sufficiently long fragment could not be obtained, fragments were additionally sequenced from the reverse direction to build a longer contig. Sequencing was performed in an ABI 3730 Genetic Analyzer (Applied Biosystems, Darmstadt, Germany). Sequences were assembled in Codon Code Aligner v5.0 (CodonCode Corporation, Dedham, Massachusetts) and ambiguities were corrected by hand. All the amplified mitochondrial sequences were classified accordingly to the standardized nomenclature of the Archie Carr Centre for Sea turtle Research (http://accstr.ufl.edu). Wright’s fixation index (F_ST_) were computed with 10.000 permutations using Arlequin v3.5.1^28^ (Electronic supplementary material, Table S3).

### Defining the geomagnetic profiles

To evaluate whether the global nesting regions have significantly different geomagnetic profiles and to test for the stability of the difference in the last century, for each nesting site, we collected geomagnetic profiles at 3 time-points (the present day, standardized as 01/08/2015, 50 and 100 years prior to the present day). This confirmation step is important since it serves as a pre-requisite for the geomagnetic field to act as a reliable source of navigation cues everywhere and on an ecologically relevant timeframe. We used the National Oceanic and Atmospheric Administration (NOAA) Magnetic Field Calculator (http://www.ngdc.noaa.gov/geomag-web/#igrfwmm) and the International Geomagnetic Reference Field (IGRF) model, by entering the GPS coordinate of each natal area. The resulting geomagnetic field vectors included: declination and inclination in degrees (°), horizontal intensity in nano-teslas (nT) as well as the northern component (nT), eastern component (nT), vertical component (nT) and total field intensity (nT) at each natal area.

### Estimating migration error rate

To estimate sea turtle navigational error (error rate), we focused on distances between isolines (bands of equal magnetic intensity) within a region. We first established the most northern and southern natal areas and their coordinates within each region. Next, we obtained geographic coordinates at each change of 1000nT (i.e. a change in 1 isoline) between these coordinates using the NOAA Magnetic Field Calculator (https://www.ngdc.noaa.gov/geomag-web/#igrfgrid) with the IGRF. Each region was given a standardized longitude comprised of the mean longitude of its most northern and southern locations. We then calculated the distance in kilometres between the geographic coordinates of each isoline using the NOAA Latitude/Longitude Distance Calculator. A region’s navigational error rate is therefore given by the mean of the distances between its isolines.

### Magnetic variation across regions

To quantify the selection acting on turtle philopatric accuracy, we also assessed how much error a turtle might accrue if it mistook adjacent vertical and total field intensity isolines. Vertical field intensity and total field intensity at each 1-degree change in latitude across regions were collected using the NOAA Magnetic Field Calculator with the IGRF. We defined each region by their most northern and most southern natal areas and longitude for a region was standardized as the mean of the longitudes of the natal areas within the region. This particular data collection was done in 2-year intervals across 24 years till the present day (01/08/1991 – 01/08/2015), based on the time it would take a turtle to sexually mature from hatching. Natal locations were categorized into northern or southern hemisphere based on their latitude.

### Data analyses

Statistical analyses were performed in R Studio version 3.2.5 [(C) 2016 The R Foundation for Statistical Computing]. Three between-class analyses (BCA from the *ade4* package (Dray & Dufour 2007)) were performed (for the present day, 50 and 100 years prior to the present day) to establish whether geomagnetic field vectors differ among sea turtle nesting regions worldwide and whether these differences were stable across time.

To test for the possible correlation between geomagnetism and population structure (indices of population genetics differentiation at specific locations), the individual geomagnetic vectors were converted into distance matrices using the maximum distance (supremum norm) between any two points of a breeding region. We used the maximum distance as this accounts for potential land-barriers and Euclidean or ordinary-line distance would oversimplify distances among natal locations. As the geomagnetic field is intrinsically linked to geographic location (confirmed here; Pearson’s product-moment correlation, t = 39.658, df = 1287, p < 0. 001), geomagnetism in our models was used as the residuals of the correlation between the magnetic field and geographic distance. Our global model included sea turtle fixation indices as the response variable with geomagnetic residuals, geographic distance between locations, sea turtle region and the two way interactions between the geomagnetic residuals and the other variables. This model was backward selected using the Akaike’s Information Criterion (Burnham & Anderson 2004)(stepAIC). Further independent linear models were performed to explore both the relationship between sea turtle population genetic structure and the geomagnetic field as well as the relationship between the rate of error in “magneto-sensing” and “non-magneto sensing” regions in the northern and southern hemispheres.

## Results and Discussion

Sea turtles require region-specific geomagnetic vectors that are stable over at least one generation if they are to use the earth’s magnetic field to conduct their philopatric migration. Hence, we studied the geomagnetic signatures over the last century (present, 50 and 100 years before present) from all regions for the five sea turtle species for which we obtained genetic data. We confirmed that each region has had a specific geomagnetic signature at any time point over the last century (Randomization Test – Present: Obs = 0.049, p = 0.001; 50 years before present (BP): Obs = 0.827, p = 0.001; 100 years BP: Obs = 0.821, p = 0.001, Fig. 2). This result is a mandatory pre-requisite for the geomagnetic field to provide informative navigational cues across nesting regions and over time.

**Fig. 2:**
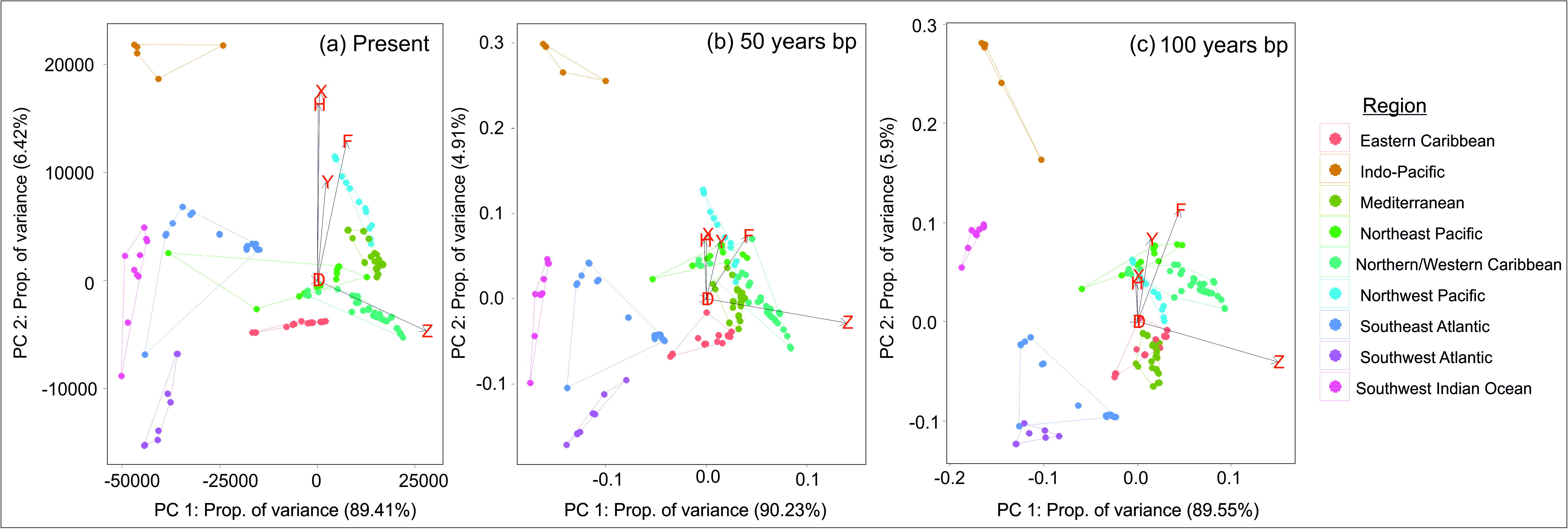
Geomagnetic fields’ representation. Principle component analysis including eigenvectors (D = Declination, I = Inclination, H = Horizontal Component, F = Total Field Intensity, X = North Component, Y = East Component, Z = Vertical Component) illustrating the geomagnetic vectors for each region from all sea turtle species. (a) The present day (01/08/2015), (b) 50 years and (c) 100 years prior to the present day.

A principal component analysis, for the present day, shows that PC1 explains 89.4% of total variance, with PC2 explaining 6.42% (50-years before present (BP) PC1 = 90.23%, PC2 = 4.91%; 100-years BP PC1 = 89.55 %, PC2 = 5.9%). For PC1, 78.6% was explained by the vertical field component (Z) followed by total field intensity (F) with 21.3%. The pattern remained the same for both 50 (Z = 76.9%, F = 23.1% of PC1) and 100 years BP (Z = 77.2%, F = 22.8% of PC1). Thus, over the last century, this vector has been stable enough for turtles to differentiate between nesting regions. Previous studies showed that sea turtles from Florida can perceive magnetic total field intensity (Lohmann & Lohmann 1996; Lohmann *et al*. 2001; Lohmann *et al*. 2008b; Putman *et al*. 2011), an element of which is the vertical field component. Ultimately, this finding shows that geomagnetism is a basis acting as a navigational cue that underlies philopatric behaviours in sea turtle species at a time-scale relevant with both ecological (navigation) and evolutionary (population structure) processes.

As a second step, we investigated the correlation between geomagnetic vectors and indices of population structure for the five sea turtle species and 144 locations (Table S1). We found that within nesting regions, the stronger the geomagnetic difference between nesting groups is, the stronger the population structure (F_1, 694_ = 52.845, p < 0.001). We also found differences across species (F_4, 715_ = 17.37, p < 0.001) with generally higher genetic structure for the Hawksbill turtle (mean ± sd F_ST_ = 0.451 ± 0.283) and lowest for the Olive Ridley (F_ST_ 0.009 ± 0.044). Strikingly, no interaction between species and geomagnetic vectors was detected (F_4, 694_ = 0.997, p = 0.409). Instead, an interaction between regions and geomagnetic vectors was a better predictor of the observed genetic structure (F_7, 694_ = 11.620, p < 0.001, Fig. 3a). Specifically, we found that the Northern/Western Caribbean, Northeast Pacific and Southwest Indian Ocean showed the expected positive correlation between genetic structure and geomagnetic field following the geomagnetic imprinting hypothesis (Northern/Western Caribbean: F_1, 256_ = 12.75, p < 0.001, Northeast Pacific: F_1, 21_ = 6.318, p = 0.020, Southwest Indian Ocean: F_1, 43_ = 3.772, p = 0.059). Crucially, this pattern was observed for all turtle species nesting within those regions, i.e. Green turtle, Hawksbill, Leatherback, Loggerhead and the Olive Ridley (Electronic supplementary material, Table S1). Thus, even though the absolute level of genetic differentiation may vary among sea turtle species, all turtles nesting in those three regions have retained parallel use of geomagnetic profile as a cue for their philopatric migration.

**Fig. 3a & b:**
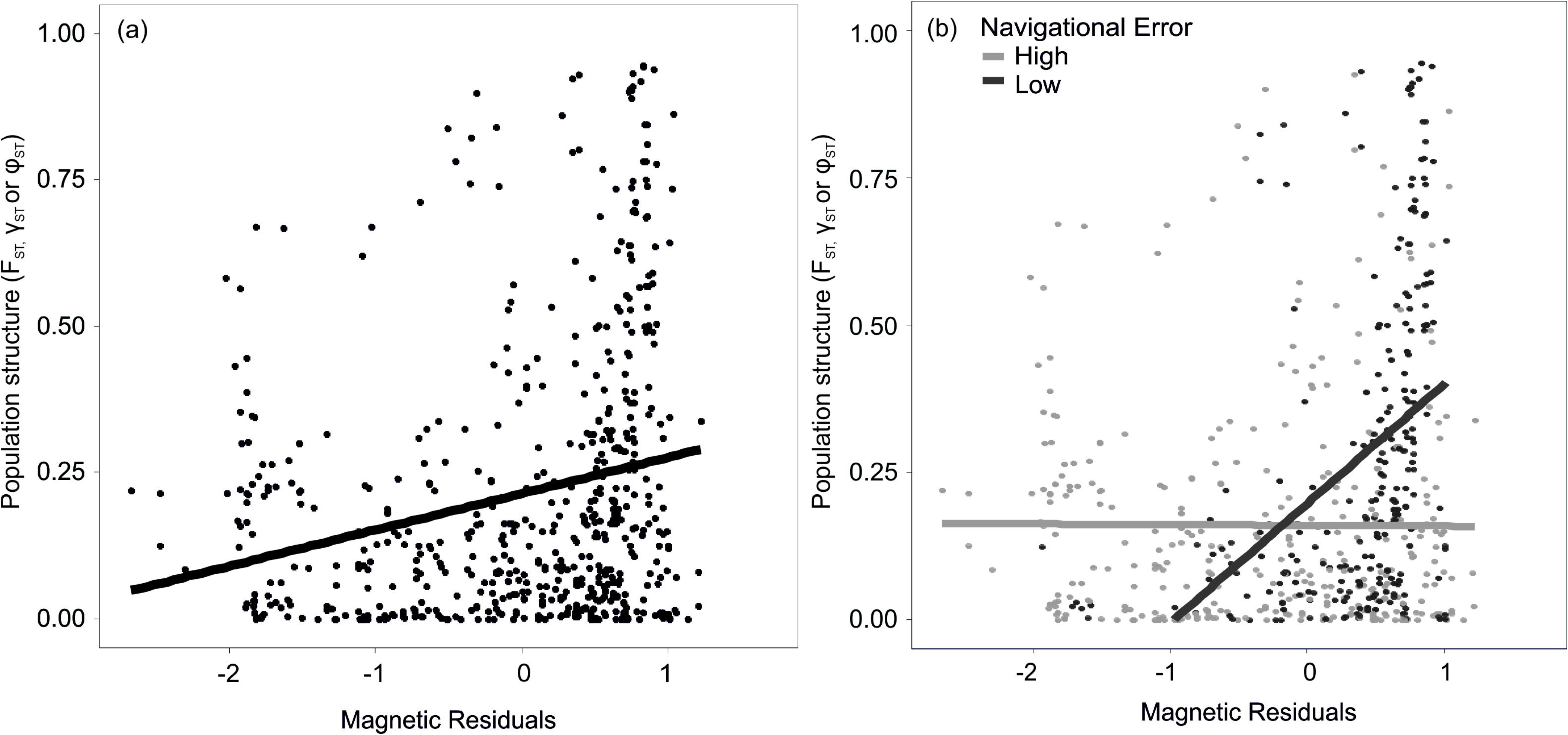
Relationship between sea turtle population structure and the geomagnetic field. The geomagnetic field is expressed as residuals of the correlation between the geomagnetic field and geographic distance. (a) Shows the overall relationship for all regions and (b) these relationships for geomagnetic regions, that offer low rates of navigational error, and for non-geomagnetic (high rates of navigational error) regions.

Conversely, in the six other nesting regions, we did not detect the expected relationship between population genetic structure and region-specific magnetic field. Here as well, this is true for all species nesting within those regions (All F < 1.199, all p > 0.281, Fig. 3b) and suggests that, in these regions, the magnetic field might not provide sufficient information to locate natal areas (i.e. area around their nesting site) with sufficiently high accuracy. Noteworthy, as expected, across the globe and across species, geographic distance was a good predictor of indices of population structure (F_1, 694_ = 64.163, p < 0.001). Overall, we demonstrate that geomagnetism-mediated philopatric behaviour is region-specific and not species-specific. This conclusion is reinforced by the parallel patterns observed across species within a region. But what are the characteristics of the earth magnetic field in some regions compared to others that would explain geomagnetic imprinting?

For each geomagnetic vector, the Earth is stratified by lines of constant magnetic values, known as isolines (Lohmann *et al*. 2008b)(Fig. 1). As coastlines are marked by isolines, sea turtles are thought to follow magnetism of equal value back to the area near their nesting sites (Lohmann *et al*. 2008b; Endres *et al*. 2016). Aggregated magnetic isolines across a geographic area highlight intense clines of the specific geomagnetic vector, whereas more dispersed isolines over the same distance represent weaker gradients of the same vector. We tested whether the different groups of regions varied in the vertical and total field intensity vectors, which we show best characterize the specific rookeries. We also assessed whether there was any consistency in the variation across both hemispheres. We found magnetic-regions, i.e. those with correlations between population structure and the geomagnetic vectors, to differ significantly from non-magnetic regions for vertical field intensity in both the northern and the southern hemisphere (Northern Hemisphere: F_1, 1376_ = 383.5, p < 0.001, Southern Hemisphere: F_1, 713_ = 219.5, p < 0.001, Fig. 4a). Magnetic regions, therefore, are those that typically have increased vertical field intensity compared to non-magnetic regions, regardless of hemisphere. Magnetic and non-magnetic regions also differ in their total field intensity in the northern hemisphere (F_1, 1376_ = 120.7, p < 0.001), however no difference was found between regions in the southern hemisphere (F_1, 713_ = 2.087, p = 0.149, Fig. 4b).

**Fig. 4a & b:**
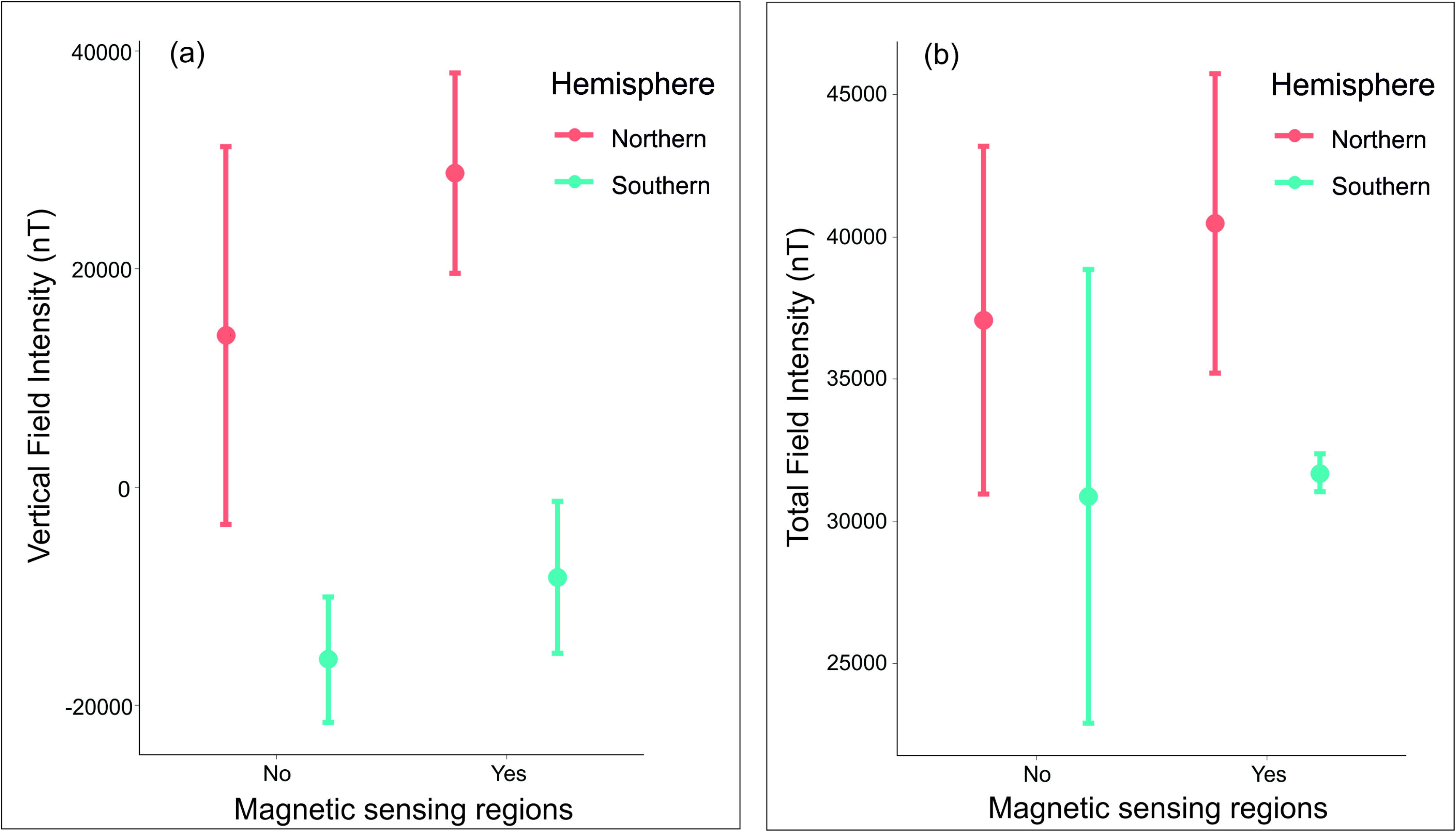
Magnetic field vector intensity for geomagnetic and non-geomagnetic regions split by hemisphere. Means and standard deviation depicted. Data collected at each change of 1 degree of latitude, across each region from the most northern sea turtle natal areas to the most southern. Each data point (isoline) was sampled over 24 years from 01/08/1991 to 01/08/2015 at 2-year intervals, i.e. the average time a sea turtle would take to mature. (a) Shows vertical field intensity and (b) shows total field intensity.

To quantify the selection acting on turtle philopatric accuracy, we determined how far a turtle might stray if it mistakenly followed adjacent vertical and total field intensity isolines. We show that sea turtles from geomagnetic regions have significantly lower rates of philopatric error (223km ± 93km) based on total field intensity compared to those in non-geomagnetic regions (390km ± 184km, F_1, 217_ = 8.366, p < 0.001, Fig. 5). In contrast, no such difference was found between geomagnetic regions (120km ± 21km) and non-geomagnetic regions (124km ± 46km) for the error-rate if sea turtles were to use vertical field isolines alone (F_1, 227_ = −0.889, p = 0.375). This result is consistent with research based on hatchling geomagnetic orientation, which follow bi-coordinate geomagnetic map from which both longitudinal and latitudinal information can be derived (Lohmann *et al*. 2008b; Putman *et al*. 2011). As the vertical field only gives information towards the magnetic equator (i.e. equator = 0), it is unlikely sea turtles would use it alone. Instead, turtles likely derive information from the combined use of vertical and total field intensity.

**Fig. 5:**
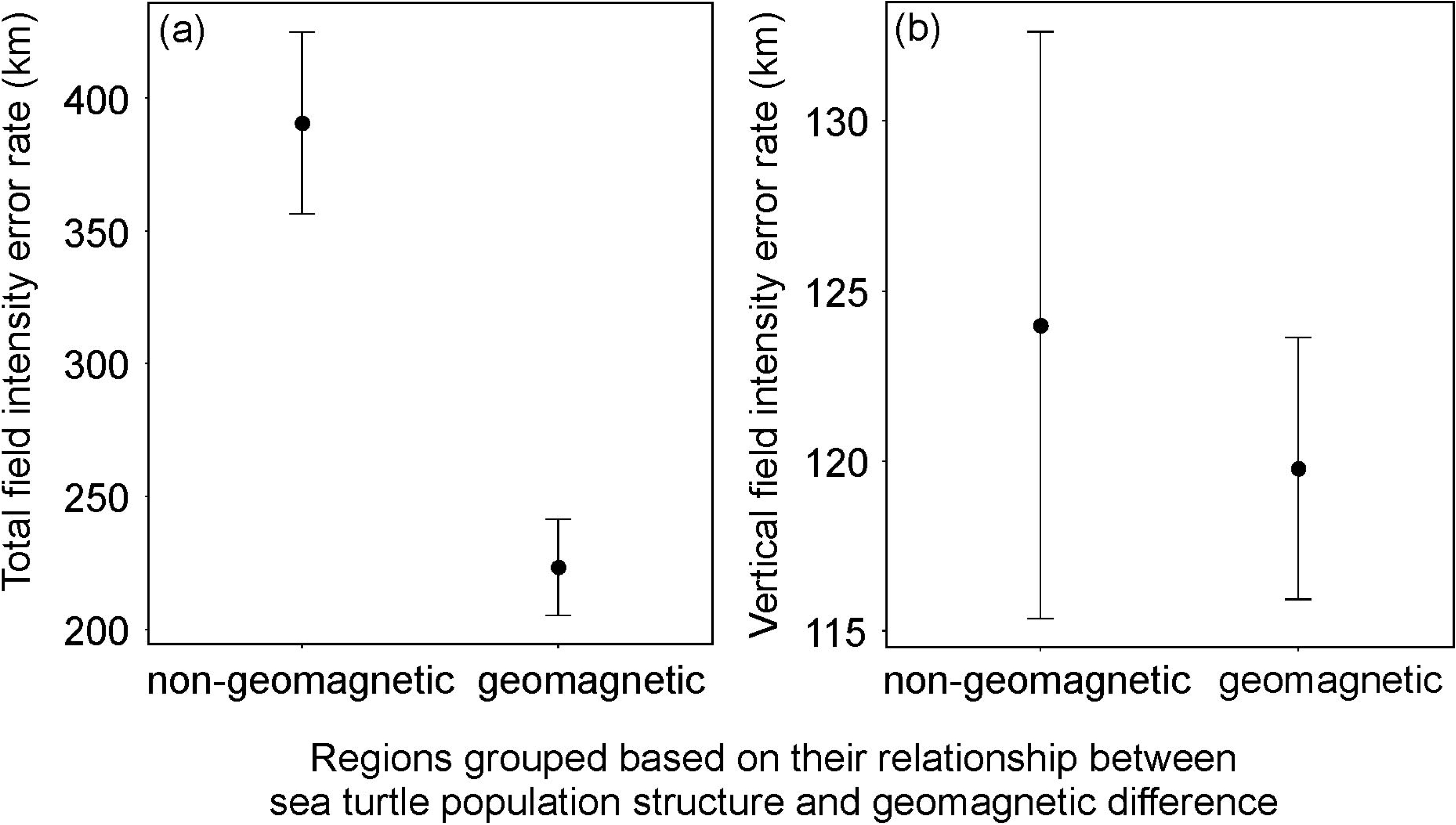
Estimated error rate in geomagnetic and non-geomagnetic regions. Means and confidence intervals are depicted. Values are obtained for “geomagnetic” and “non-geomagnetic” regions from the estimation of distance if turtles confused two adjacent isolines for magnetic total field intensity and vertical field intensity.

While our findings are consistent with previous studies (Lohmann *et al*. 2008b; Endres *et al*. 2016; Brothers & Lohmann 2018), we demonstrate that geomagnetic imprinting is regional, not universal. This result concurs with secular variation of the Earth’s magnetic field, which causes female sea turtles to change natal locations as magnetic signatures drift along coastlines (Brothers & Lohmann 2015). In general, our result demonstrates that slight inaccuracy in the detection of the geomagnetic field vector may lead turtles to fail to return to breeding sites (Lohmann *et al*. 2008b) creating extreme selection pressure to increase nesting site fidelity.

Overall, our study reveals the selective pressure exerted by the geomagnetic field on sea turtle population structure and how this contributes to sea turtle evolution. In geomagnetic-regions, isolines are close enough to one another to provide sufficiently accurate navigational information for turtles to reach their natal area and reproduce. This pattern of philopatry will ultimately restrict gene flow among populations within regions and among regions. In regions with distant isolines, navigation mistakes could result in the best case in increased gene flow among natal areas, or in the worst case in failure to reproduce at all. In these regions, philopatric journeys will likely require sea turtles to utilize multi-modal cues, e.g. olfaction and geomagnetism (Lohmann *et al*. 1999; Lohmann *et al*. 2008a; Endres *et al*. 2016). Previous genetic analyses also show that the accuracy of sea turtle philopatric journeys vary between different populations (Meylan *et al*. 1990; Bowen & Karl 2007; Lee *et al*. 2007) and our results could suggest these differences in accuracy may be determined by local cue. It would also explain why turtles with disrupted magneto-sensing capacity by the application of magnet on their head, despite longer journey, find their natal area (Luschi *et al*. 2007).

In our study, we cannot account for all the other parameters that also influence the distribution of sea turtles. For instance, modern and historical climates (Poloczanska *et al*. 2009; Pike 2013), ocean currents (Putman *et al*. 2010) or plate tectonics (Bowen *et al*. 1989) have all been implicated in the distribution of sea turtles. Our results suggest that rookery-specific environmental pressures will define which parameter is the most important. Interestingly, the framework we have detailed in this study should facilitate testing the different possible drivers of population genetics and ultimately determine their relative contribution in other species that have evolved philopatric behaviours.

In conclusion, while our results demonstrate that there is sufficient differentiation among the geomagnetic fields of sea turtle rookeries, location-specific use of given vectors suggests the parallel (across species) and local (to regions) adaptation of navigational cue recognition. Whilst our analyses focus specifically on sea turtles, its relevance extends to other taxa conducting long-distance migrations to and from their natal areas. Many other marine organisms e.g. elephant seals, European eels or salmon have been hypothesized to imprint on a natal geomagnetic field to complete their philopatric migrations (Matsumura *et al*. 2011; Putman *et al*. 2013; Putman *et al*. 2014; Baltazar-Soares & Eizaguirre 2017; Naisbett-Jones *et al*. 2017). Our results suggest that underlying mechanisms for geomagnetic navigation are locally adapted and future research into such mechanisms should focus on evolved region-specific and coupled systems rather than a universal molecular/physiological mechanism.

## Acknowledgements

We would like to thank the members of the Eizaguirre Lab for discussions and comments on earlier versions of the manuscript. We also would like to thank G. Schofield for comments on a previous version of this manuscript.

## Author Contributions

CE and SJKC designed the study, CE and SJKC conducted sampling in Cape Verde. CE and MBS conducted the molecular analyses. SJKC performed all the statistics of the meta-analysis and drafted the manuscript. All authors edited the manuscript and contributed to its final version.

