## Supplementary Material for "Region-specific magnetic fields structure sea turtle populations"

**Electronic supplementary material, Table S1:** List of literature utilized within this study with corresponding species and assigned region

| Species | Region | Literature |
| --- | --- | --- |
| Green | Mediterranean | Bagda et al 2012 |
| Green | Southwest Indian Ocean | Bourjea et al 2007 |
| Green | Northeast Pacific | Chassin-Noria et al. 2004 |
| Green | Mediterranean | Costa-Jordao 2017 |
| Green | Northern/Western Caribbean | Costa-Jordao 2017 |
| Green | Eastern Caribbean | Costa-Jordao 2017 |
| Green | Southeast Atlantic | Costa-Jordao 2017 |
| Green | Southwest Atlantic | Costa-Jordao 2017 |
| Green | Southeast Atlantic | Formia et al 2006 |
| Green | Northern/Western Caribbean | Naro-Marciel et al 2014 |
| Green | Eastern Caribbean | Naro-Marciel et al 2014 |
| Green | Southwest Atlantic | Naro-Marciel et al 2014 |
| Green | Northwest Pacific | Nishizawa et al 2011 |
| Green | Northern/Western Caribbean | Ruiz-Urquiola et al, 2010 |
| Green | Northern/Western Caribbean | Shamblin et al 2015 |
| Green | Eastern Caribbean | Shamblin et al 2015 |
| Hawksbill | Northeast Pacific | Gaos et al, 2016 |
| Hawksbill | Northern/Western Caribbean | Leroux et al, 2012 |
| Hawksbill | Eastern Caribbean | Leroux et al, 2012 |
| Hawksbill | Northern/Western Caribbean | Monzon-Arguello et al 2011 |
| Hawksbill | Eastern Caribbean | Monzon-Arguello et al 2011 |
| Hawksbill | Northern/Western Caribbean | Proietti et al, 2014 |
| Hawksbill | Eastern Caribbean | Proietti et al, 2014 |
| Leatherback | Indo-Pacific | Dutton et al 1999 |
| Leatherback | Eastern Caribbean | Dutton et al 1999 |
| Leatherback | Northern/Western Caribbean | Dutton et al 1999 |
| Leatherback | Northeast Pacific | Dutton et al 1999 |
| Leatherback | Southeast Atlantic | Dutton et al 2013 |
| Loggerhead | Northern/Western Caribbean | Bowen et al 2005 |
| Loggerhead | Mediterranean | Carreras et al 2007 |
| Loggerhead | Mediterranean | Clusa et al 2013 |
| Loggerhead | Mediterranean | Garofalo et al 2009 |
| Loggerhead | Northwest Pacific | Hatase et al 2002 |
| Loggerhead | Southeast Atlantic | Klein, Balthazar-Soares et al (in prep) |
| Loggerhead | Northwest Pacific | Matsuzawa et al 2016 |
| Loggerhead | Southwest Atlantic | Reis et al 2010 |
| Loggerhead | Mediterranean | Saied et al 2012 |
| Loggerhead | Mediterranean | Saied et al 2012 |
| Loggerhead | Northern/Western Caribbean | Shamblin et al 2012 |
| Loggerhead | Northern/Western Caribbean | Shamblin et al 2014 |
| Loggerhead | Mediterranean | Shamblin et al 2014 |
| Loggerhead | Southeast Atlantic | Shamblin et al 2014 |
| Loggerhead | Southwest Atlantic | Shamblin et al 2014 |
| Loggerhead | Mediterranean | Yilmaz et al 2011 |
| Olive Ridley | Northeast Pacific | Lopez-Castro & Rocha-Olivares, 2005 |
| Olive Ridley | Indo-Pacific | Jensen et al 2013 |

**Electronic supplementary material, Table S2: Regions, and the 144 individual natal areas and nesting sites (locations) that characterize them, analysed in this study. Locations based on those with genetic structure information from literature.**

| Location | Region |
| --- | --- |
| Akamas, Cyprus | Mediterranean |
| Akyatan, Turkey | Mediterranean |
| Alagadi, Cyprus | Mediterranean |
| Alata Mahallesi, Turkey | Mediterranean |
| Aldabra, Seychelles | Southwest Indian Ocean |
| Amami, Japan | Northwest Pacific |
| Anamur, Turkey | Mediterranean |
| Antigua (Jumby Bay) | Eastern Caribbean |
| Arenas Blancas, Panama | Northeast Pacific |
| Ascension Island | Southeast Atlantic |
| Aves Island, Venezuela | Eastern Caribbean |
| Bahia de Jiquilisco, El Salvador | Northeast Pacific |
| Bahia, Brazil | Southwest Atlantic |
| Barbados-Leeward | Eastern Caribbean |
| Barbados-Windward | Eastern Caribbean |
| Belize (Gale's Point) | Northern/Western Caribbean |
| Bioko, Equatorial Guinea | Southeast Atlantic |
| Boa Vista, Cape Verde | Southeast Atlantic |
| Boca Raton, Florida, USA | Northern/Western Caribbean |
| Bōsō, Japan | Northwest Pacific |
| Buck Island (St Croix) | Eastern Caribbean |
| Calabria, Italy | Mediterranean |
| Canaveral National Seashore, Florida, USA | Northern/Western Caribbean |
| Cape Island, South Carolina, USA | Northern/Western Caribbean |
| Cape San Blas, Florida, USA | Northern/Western Caribbean |
| Casey Key, Florida, USA | Northern/Western Caribbean |
| Cay Sal Bank, Bahamas | Northern/Western Caribbean |
| Chiapas, Mexico | Northeast Pacific |
| Colola, Mexico | Northeast Pacific |
| Cosmoledo, Seychelles | Southwest Indian Ocean |
| Costa Rica (Playa Grande) | Northern/Western Caribbean |
| Costa Rica (Tortuguero) | Northern/Western Caribbean |
| Costa Rica (Pacific) | Northeast Pacific |
| Crete | Mediterranean |
| Cuba (Doce Leguas) | Northern/Western Caribbean |
| Cyprus | Mediterranean |
| Dalaman, Turkey | Mediterranean |
| Dalyan, Turkey | Mediterranean |
| Drt Tortugas, Florida, USA | Northern/Western Caribbean |
| E Ishigaki Island, Japan | Northwest Pacific |
| El Mansouri, Lebanon | Mediterranean |
| Enshu-nada, Japan | Northwest Pacific |
| Espírito Santo, Brazil | Southwest Atlantic |
| Estero Padre Ramos | Northeast Pacific |
| Europa island | Southwest Indian Ocean |
| Farquhar Atoll | Southwest Indian Ocean |
| Fethiye, Turkey  Flinders Beach | Mediterranean  Indo-Pacific |
| Fogo, Cape Verde | Southeast Atlantic |
| French Guinana | Eastern Caribbean |
| Ft. Lauderdale, Florida, USA | Northern/Western Caribbean |
| Fukiagehama, Japan | Northwest Pacific |
| Gabon (Mayumba beach) | Southeast Atlantic |
| Georgia, USA | Northern/Western Caribbean |
| Ghana (Ade Foah Beach) | Southeast Atlantic |
| Glorieuses | Southwest Indian Ocean |
| Guadeloupe | Eastern Caribbean |
| Guanahacabibes (Cuba) | Northern/Western Caribbean |
| Guerrero, Mexico | Northeast Pacific |
| Hillsboro, Pompano, Ft. Lauderdale, USA | Northern/Western Caribbean |
| Hutchinson Island, Florida, USA | Northern/Western Caribbean |
| Isla Cozumel, Quintana Roo, Mexico | Northern/Western Caribbean |
| Isla de Aves, Venezuela | Eastern Caribbean |
| Israel | Mediterranean |
| Juan de Nova island | Southwest Indian Ocean |
| Juno Beach, Florida, USA | Northern/Western Caribbean |
| Kazanli, Turkey | Mediterranean |
| Keewaydin Island, Florida, USA | Northern/Western Caribbean |
| Kuriat Islands, Tunisia | Mediterranean |
| Kyparissia, Greece | Mediterranean |
| Lakonikos, Greece | Mediterranean |
| Lara Bay, Cyprus | Mediterranean |
| Lebanon | Mediterranean |
| Libya | Mediterranean |
| Machalilla, Ecuador | Northeast Pacific |
| Maio, Cape Verde | Southeast Atlantic |
| Terengganu, Malaysia | Indo-Pacific |
| Marquesas Keys, Florida, USA | Northern/Western Caribbean |
| Maruata, Mexico | Northeast Pacific |
| Matapica, Suriname | Eastern Caribbean |
| Mayotte island | Southwest Indian Ocean |
| Melbourne Beach, Florida, USA | Northern/Western Caribbean |
| Isla Holbox, Quintana Roo, Mexico | Northern/Western Caribbean |
| Playa Mexiquillo, Mexico | Northeast Pacific |
| Minabe, Japan  McCluer Group | Northwest Pacific  Indo Pacific |
| Misurata, western Libya | Mediterranean |
| Miyazaki, Japan | Northwest Pacific |
| Moheli, Comoros | Southwest Indian Ocean |
| N.Cyprus | Mediterranean |
| Nicaragua (Pearl Cays) | Northern/Western Caribbean |
| North Carolina, USA | Northern/Western Caribbean |
| Nosy Iranja, Madagascar | Southwest Indian Ocean |
| NW Ishigaki Island, Japan | Northwest Pacific |
| Oaxaca, Mexico | Northeast Pacific |
| Okinawa Islands, Japan | Northwest Pacific |
| Okinoerabu, Japan | Northwest Pacific |
| Osa Peninsula, Costa Rica | Northeast Pacific |
| Ossabaw Island, Georgia, USA | Northern/Western Caribbean |
| Pailoa, Guinea Bissau | Southeast Atlantic |
| Paso de Noria, Mexico | Northeast Pacific |
| Principe island | Southeast Atlantic |
| Puerto Rico (Mona Island) | Eastern Caribbean |
| Rethymno, Crete | Mediterranean |
| Rio de Janeiro, Brazil | Southwest Atlantic |
| Rocas Atoll, Brazil | Southwest Atlantic |
| S.Luzia, Cape Verde | Southeast Atlantic |
| S.Nicholau, Cape Verde | Southeast Atlantic |
| S.Vicente, Cape Verde | Southeast Atlantic |
| Sal, Cape Verde | Southeast Atlantic |
| Samandağ/Hatay, Turkey | Mediterranean |
| San Felipe, Cuba | Northern/Western Caribbean |
| Santa Luzia, Cape Verde | Southeast Atlantic |
| Santiago, Cape Verde | Southeast Atlantic |
| Santo Antao, Cape Verde | Southeast Atlantic |
| Sao Tome | Southeast Atlantic |
| Sergipe, Brazil | Southwest Atlantic |
| Shikoku, Japan | Northwest Pacific |
| Sinaloa, Mexico | Northeast Pacific |
| Singer Island, Florida, USA | Northern/Western Caribbean |
| Sirte, western Libya | Mediterranean |
| Solomon Islands | Indo-Pacific |
| South Carolina, USA | Northern/Western Caribbean |
| Southwestern Cuba | Northern/Western Caribbean |
| St. George Island, Florida, USA | Northern/Western Caribbean |
| Sandy Point, St.Croix | Eastern Caribbean |
| Matapica, Suriname | Eastern Caribbean |
| SW Iriomote Island, Japan | Northwest Pacific |
| Tekirova, Turkey | Mediterranean |
| Tequesta, Southern Jupiter Island  Tiwi Island | Northern/Western Caribbean  Indo-Pacific |
| Tortuguero, Costa Rico | Northern/Western Caribbean |
| Grande Riviere beach, Trindad | Eastern Caribbean |
| Trindade Island, Brazil | Southwest Atlantic |
| Tromelin island | Southwest Indian Ocean |
| Tunisa | Mediterranean |
| Turkey | Mediterranean |
| U.S. Virgin Islands | Eastern Caribbean |
| Venezuela | Eastern Caribbean |
| Volusia County, Florida, USA | Northern/Western Caribbean |
| W. Turkey | Mediterranean |
| Yakushima, Japan | Northwest Pacific |
| Yumurtalik, Turkey | Mediterranean |
| Zakynthos Island, Greece | Mediterranean |

|  | Boavista | Fogo | Maio | Santo Antão | Sal | S.Nicolau | S.Luzia | S.Vicente | Santiago |
| --- | --- | --- | --- | --- | --- | --- | --- | --- | --- |
| Boavista | NA | 0.225 | 0 | 0.004 | 0.007 | 0.007 | 0.005 | 0.2 | 0 |
| Fogo | - | NA | 0.156 | 0.145 | 0.114 | 0.058 | 0.171 | 0.124 | 0.098 |
| Maio | - | - | NA | 0.001 | 0.01 | 0.011 | 0.006 | 0.15 | 0 |
| Santo Antão | - | - | - | NA | 0.013 | 0.018 | 0.034 | 0.137 | 0 |
| Sal | - | - | - | - | NA | 0 | 0.009 | 0.078 | 0 |
| S.Nicholau | - | - | - | - | - | NA | 0 | 0.066 | 0.006 |
| S.Luzia | - | - | - | - | - | - | NA | 0.145 | 0.011 |
| S.Vicente | - | - | - | - | - | - | - | NA | 0.082 |
| Santiago | - | - | - | - | - | - | - | - | NA |

**Electronic supplementary material, Table S3:** **Above the diagonal are F_ST_ values for nesting groups within the Cape Verde archipelago.**
